# Automated nuclear cartography reveals conserved sperm chromosome territory localization across 2 million years of mouse evolution

**DOI:** 10.1101/508770

**Authors:** Benjamin Matthew Skinner, Joanne Bacon, Claudia Cattoni Rathje, Erica Lee Larson, Emily Emiko Konishi Kopania, Jeffrey Martin Good, Nabeel Ahmed Affara, Peter James Ivor Ellis

## Abstract

Measurements of nuclear organization in asymmetric nuclei in 2D images have traditionally been manual. This is exemplified by attempts to measure chromosome position in sperm samples, typically by dividing the nucleus into zones, and manually scoring which zone a FISH signal lies in. This is time consuming, limiting the number of nuclei that can be analyzed, and prone to subjectivity. We have developed a new approach for automated mapping of FISH signals in asymmetric nuclei, integrated into an existing image analysis tool for nuclear morphology. Automatic landmark detection defines equivalent structural regions in each nucleus, then dynamic warping of the FISH images to a common shape allows us to generate a composite of the signal within the entire cell population. Using this approach, we mapped the positions of the sex chromosomes and two autosomes in three mouse lineages (Mus *musculus domesticus, Mus musculus musculus* and *Mus spretus*). We found that in all three, chromosomes 11 and 19 tend to interact with each other, but are shielded from interactions with the sex chromosomes. This organization is conserved across 2 million years of mouse evolution.

## Introduction

Studies of the sub-nuclear localisation of chromatin often use fluorescence in-situ hybridisation to detect DNA or RNA, or immunostaining to detect proteins. The images are subsequently analysed either manually, or using some automated analysis tool. If the nucleus is circular or elliptical, it is commonly divided into concentric shells of equal area and the proportion of signal in each shell is measured (e.g. [1–3]). This has been amenable to automation, allowing analysis of thousands of cells, which, with appropriate statistical treatment, can yield valuable data at a scale that is still beyond the scope of 3D imaging techniques in time and cost.

However, if the nucleus is asymmetric, such as in sperm, a shell analysis is not sufficient. Frequently, nuclei are manually divided into geometric regions, and the number of nuclei with signals in each region are counted. For example, in spatulate sperm such as pig or human, positions of loci are located into anterior, medial and posterior regions [4–6], or measured by proportional position along each axis [7]. Rodent sperm have a more interesting, falciform, hooked shape: they have two axes of asymmetry, the anterior-posterior and the dorsal-ventral axis. This means that the location of a FISH signal can - in principle - be unambiguously localised and compared between nuclei. The determination of chromosome position is still manual, with more regions of the nucleus into which a signal may be assigned [8,9], or described without quantitation [10]. This is both time-consuming, and subjective, limiting the numbers of nuclei that can be analysed.

The positions of chromosomes or other loci in gametes (particularly sperm) is of great interest due to both the association of nuclear organisation with fertility in the clinic, in agriculture, and in evolutionary biology. Chromosome position has been linked with infertility in human males; men presenting with fertility problems have less consistent chromosome territories than healthy men [11–13]. Similarly, in farm animals, studies of nuclear organisation have discovered conserved sperm chromosome territories in boars [4], and wider evolutionary studies have shown conservation of some chromosomes - such as the X - from eutherian mammals to marsupial mammals and monotremes [14].

Newer sequencing-based approaches, such as Hi-C are being used to produce 3D maps of chromatin structure across multiple - and even single - nuclei [15–17]. Validating these results by microscopy is harder due to the number of cells that must be analysed, yet is necessary for our understanding of how chromatin patterns seen across millions of cells relate to chromatin structure within an individual nucleus. Three-dimensional imaging such as confocal microscopy provides high quality position information, but is time-consuming and costly in comparison to 2D fluorescence imaging.

Given this, there is a need to quickly and robustly assay nuclear organisation in 2D fluorescence microscopy images with greater precision than is currently available. Here, we demonstrate the use of automatic landmark detection in nuclei to rapidly localise, aggregate and compare nuclear signals without need for precise detection of the signal boundaries, or extensive manual thresholding and curation. We use this method to investigate the conservation of nuclear organisation between three mouse lineages, *Mus musculus musculus, Mus musculus domesticus* and *Mus spretus*. Of these, *M. spretus* has a notably different nuclear shape [18] to the others, being shorter and wider, allowing us to test whether chromosome position is conserved across structurally equivalent regions.

## Materials and Methods

### Sample collection

We collected sperm from wild-derived inbred mouse strains *Mus musculus musculus* (PWK/PhJ), *M. m. domesticus* (LEWES/EiJ) and *Mus spretus* (STF). All animal procedures were in accordance with the University of Montana Institute for Animal Care and Use Committee (protocol 002-13) and were subject to local ethical review. Animals were bred at the University of Montana from mice purchased from Jackson Laboratories (Bar Harbor, ME) or were acquired from Francois Bonhomme (University of Montpellier). Animals were housed singly or in small groups, sacrificed via CO2 followed by cervical dislocation, and tissues were collected *post mortem* for analysis. Sperm were collected and fixed in 3:1 methanol-acetic acid as previously described [18].

### Fluorescence in-situ hybridisation (FISH)

Fixed sperm were dropped on poly-lysine slides, air-dried, and aged at 70°C for one hour. Sperm were swelled in 10mM DTT in 0.1M Tris-Hcl for 30 minutes at room temperature (RT). Slides were rinsed in 2xSSC (saline sodium citrate) and dehydrated through an ethanol series (70%, 80%, 100%, 2mins at RT). Chromatin was relaxed by incubating slides in 0.1mg/ml pepsin in 0.01N HCl at 37°C for 20 mins. Nuclei were permeabilized in 0.5% IGEPAL CA-630, 0.5% Triton-X-100 at 4°C for 30 minutes, and dehydrated through an ethanol series. Slides and chromosome paints for chrX, Y, 11 and 19 (Cytocell, AMP-0XG, AMP-0YR, AMP-11G, AMP-19R) were separately denatured in 70% formamide at 75°C for 5 minutes, then slides were dehydrated through an ethanol series. Probes were co-hybridised in pairs of 4μl each of: chrX and chrY; chrX and chr19; chr11 and chr19. The probes were added to the slides, coverslips were sealed with rubber cement, and the slides were hybridised for 48 hours at 37°C. Coverslips were removed, and slides were washed in 0.7xSSC, 0.3% Tween-20 at 73°C for 3 minutes to remove unbound probe, then washed in 2xSSC for 2 minutes at RT, rinsed in water and air-dried in the dark. Slides were counterstained with 16μl VectorShield with DAPI (Vector Labs) under a 22×50mm cover slip and imaged at 100x on an Olympus BX-61 epifluorescence microscope equipped with a Hamamatsu Orca-ER C4742-80 cooled CCD camera and appropriate filters. Images were captured using Smart-Capture 3 (Digital Scientific UK) with fixed exposure times for each fluorochrome.

### Image analysis

Analysis was performed using our image analysis software (Nuclear Morphology Analysis, available from http://bitbucket.org/bmskinner/nuclear_morphology/wiki/Home/, version 1.15.0) for morphometric analysis of mouse sperm shape [18]. Here, we combine nuclear morphometry with FISH signal detection in order to rigorously quantify the distribution of chromosome territories within the asymmetric mouse sperm head. Within our images we detected 1445 PWK nuclei, 906 LEWES nuclei and 712 STF nuclei across all hybridisations (Figure 1B). The number of nuclei with FISH signals detected which were used for chromosome positioning analysis are given in Supplementary Table 1.

**Figure 1.**
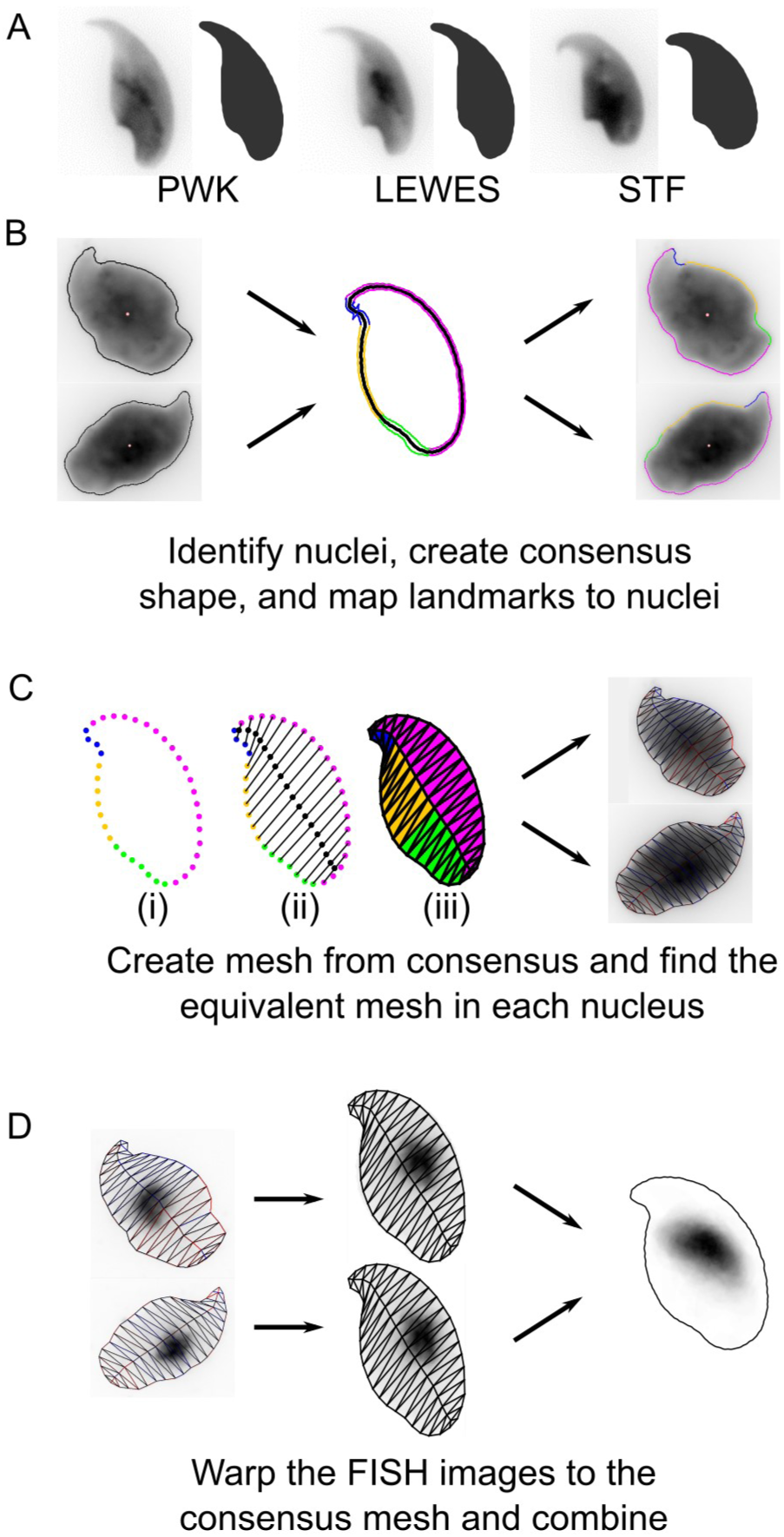
The process of warping FISH images. A) Examples of un-FISHed nuclei from the three strains, as described in [18]. B) After FISH, nuclei are automatically identified and landmarks are discovered. C) A mesh is created from the consensus nuclear shape; (i) peripheral vertices are evenly spaced between landmarks; (ii) internal vertices divide vertex pairs from the tip; (iii) all vertices are joined. The equivalent mesh is constructed for each nucleus. D) The FISH signal image is transformed to move every pixel to its location in the consensus mesh. The warped images are combined to yield the composite signal image.

This analysis, which we refer to as “nuclear cartography” is a form of mesh warping, achieved by overlaying a mesh onto each individual sperm nucleus and quantifying the distribution of the chromosomal signal within each face of the mesh (Figure 1C). This allows accurate, quantifiable 2D analysis of the signal distribution in each cell. Subsequently, since the mesh overlaid onto each sperm head is structurally equivalent, dynamic image warping is used to combine multiple individual nuclear outlines onto the consensus shape of the cell population (Figure 1D). Using this method, signal intensity can be averaged over multiple sperm heads, reducing the effect of background inhomogeneities and revealing the consensus two-dimensional location of the signal in the population as a whole.

For successful warping of the source image, the face of the mesh to which each pixel belongs must be determined. The critical step is the construction of the mesh, such that each face contains a structurally equivalent region of the nucleus. First, we identify key landmarks around the periphery of the nucleus (i.e. the apical hook, tail attachment site, and other areas of maximal curvature), as described previously [18]. Next, semi-landmarks are constructed by spacing a set number of equidistant points between each landmark (Figure 1Ci). These then serve as the peripheral vertices of the mesh. The internal vertices are created by walking through the points pairwise from the tip of the nucleus, and generating a vertex at the centre of the line connecting each pair (Figure 1Cii). Internal and peripheral vertices are connected into the faces of the mesh (Figure 1Ciii). The same structural mesh is created for the consensus nucleus shape, and for each individual nucleus. An affine transform is applied to image pixels within each face, moving them to their equivalent positions in the consensus mesh. After pixels have been relocated, a gap-filling kernel sets any empty pixel to the average of the surrounding non-zero 8-connected pixels, as long as there are at least 4 non-zero surrounding pixels. This reduces ‘smearing’ in cases where there is a large size difference between source and consensus mesh faces.

In this way, we ‘warp’ the original images to fit the consensus nucleus. The warped images can be combined to reveal the locations of consistent nuclear signal. Random noise is averaged out, while consistent signals are reinforced. To avoid bias from higher or lower intensity signals in different nuclei, the FISH images are binarised before warping. Since the individual images are being warped to fit a template shape, it is possible to choose any template with the same underlying graph structure in the mesh. This allows comparison of FISH signal distributions between different hybridisations.

To compare signal distributions between warped signals, we used an open source implementation of a multi-scale structural similarity index measure, MS-SSIM* [19,20], which quantifies visual similarity between images [21]. To further assess co-localization, we identified the chromosomal signals within the nuclei by thresholding [3], and measured the distances between the centres of mass of co-hybridised chromosomes. Statistical analyses were performed in R 3.5.1 [22], and charts were generated using the cividis colour palette [23].

## Results

This section may be divided by subheadings. It should provide a concise and precise description of the experimental results, their interpretation as well as the experimental conclusions that can be drawn.

### The sex chromosomes have conserved position in mouse sperm nuclei

The process of hybridising FISH probes to sperm nuclei required a considerable swelling step due to the highly compact chromatin. This swelling distorts the nuclear shape; our method for automated nucleus and landmark detection [18] was able to identify and orient swelled nuclei successfully, despite the fewer landmarks available.

Confident that we could orient a FISH signal within the nucleus, we applied the new technique to FISH images of mouse sperm from three strains, using chromosome paints for the X and Y chromosomes. These have been previously reported in C57Bl6 to lie under the acrosome [8,9]. Nuclei and signals were detected from the captured images, a consensus nuclear shape was calculated for each strain, and each FISH image was warped onto that consensus shape. A composite image was created by layering each FISH image, providing - effectively - a heat-map of signal location within the nucleus.

Our results confirm a consistent sub-acrosomal location for both X and Y chromosomes (Figure 2). Following the signal warping onto the population consensus, we observed that both X and Y chromosomes have overlapping territories (Figure 3, 4).

**Figure 2.**
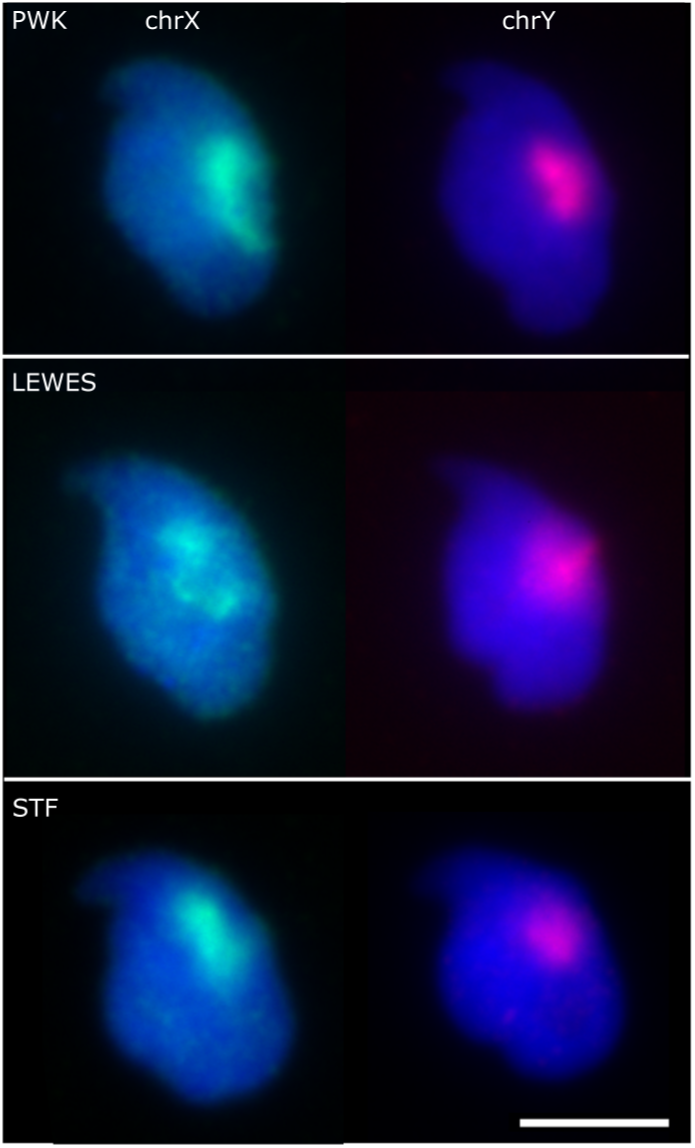
Example images showing the sex chromosome positions within the three strains. Scale bar represents 5μm.

**Figure 3.**
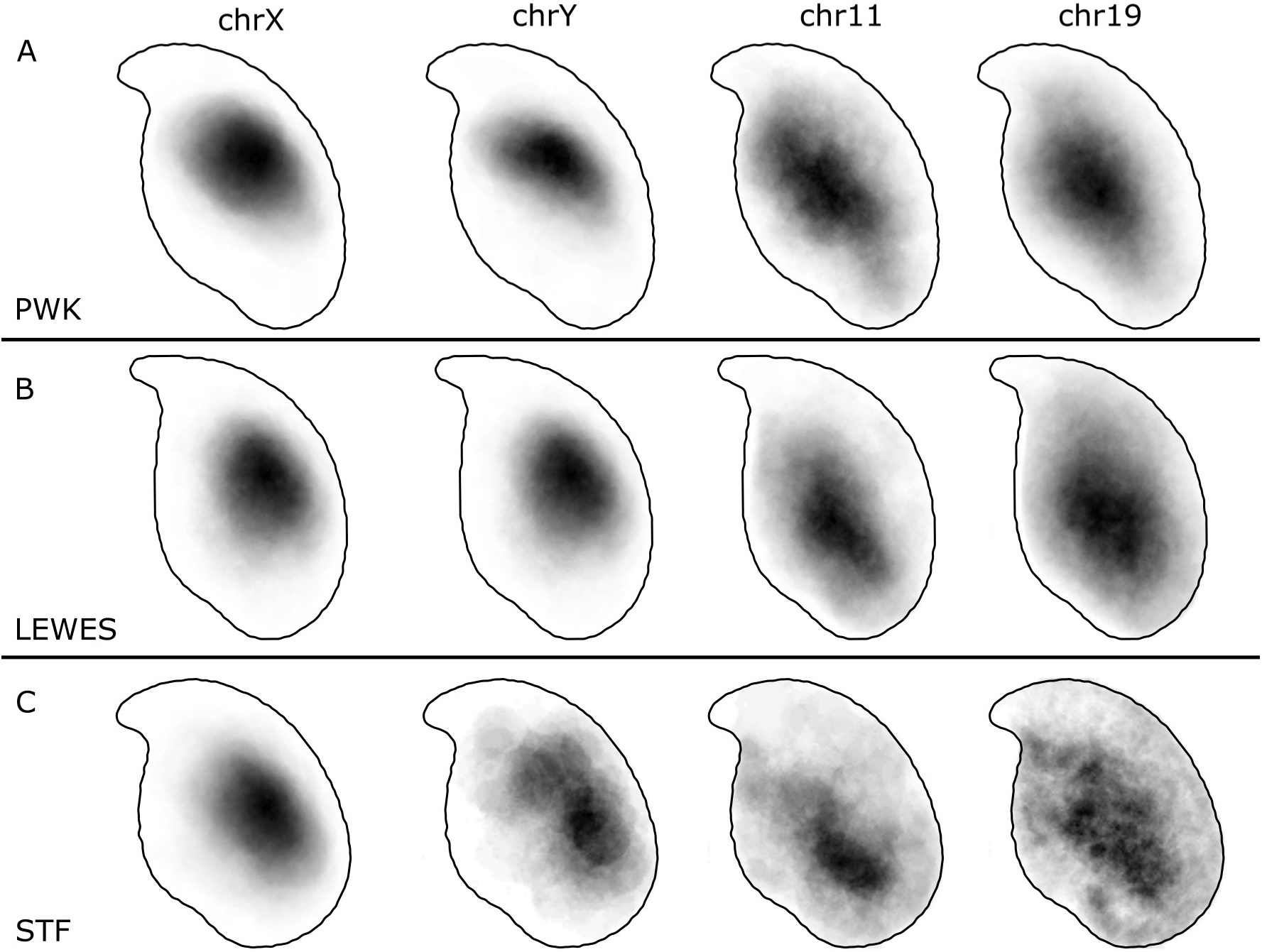
Composite signal distributions for chromosomes X, Y, 11 and 19 in (A) PWK, (B) LEWES and (C) STF. The sex chromosomes occupy a consistent territory apical and dorsal to the centre of mass, generally under the acrosome but rarely extending fully to the periphery of the nucleus. Chromosomes 11 and 19 are more widely distributed, with the predominant location basal and ventral to the centre of mass.

**Figure 4.**
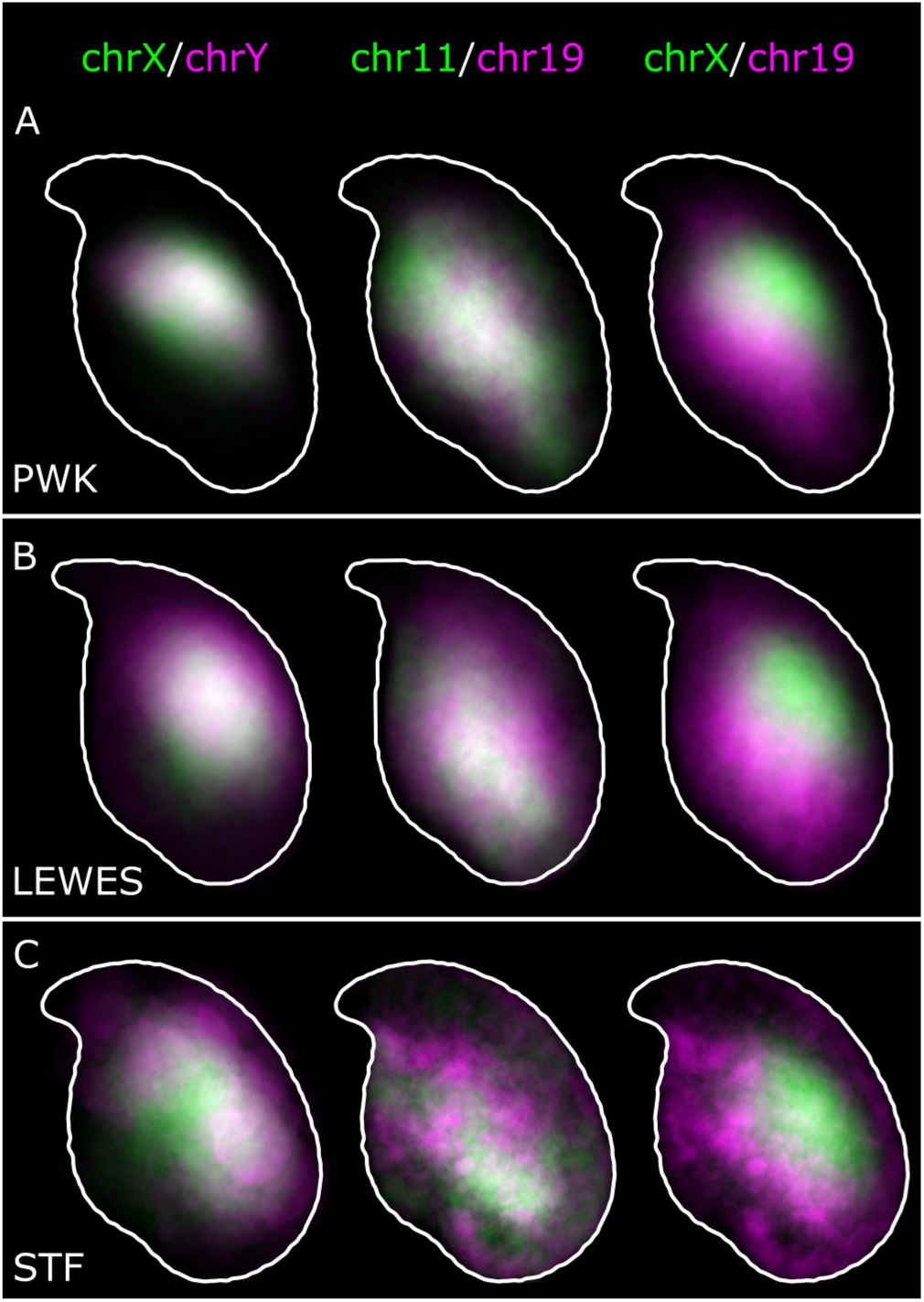
Overlay of warped distributions from Figure 3 shows the similarities between chromosome X and Y territories, and 11 and 19 territories in (A) *M. m. musculus*; (B) *M. m. domesticus*; and (C) *M. spretus*. Chromosomes X and 19 (and X and 11) are predominantly non-overlapping.

### Chromosomes 11 and 19 occupy similar nuclear addresses

With the sex chromosome locations confirmed to be conserved, we decided to examine two further chromosomes, both of which have previously been reported in the literature. Chromosome 19 has been described in C57Bl/6 mice to frequently lie toward the base of the nucleus [8]. Furthermore in Hi-C experiments, chromosomes X and 19 had a low association in C57BL sperm chromatin; chromosome 19 and chromosome 11 had a moderate association with each other [17]. For this reason, we hypothesised that chr11 and chr19 might share a similar distribution, and that this would be distinct from that of the sex chromosomes.

The composite signal position data are shown in Figure 3. The patterns are indeed different to that of the sex chromosomes. The majority of the signal lies ventral and basal to the centre of the nucleus, yet there are clearly instances of signal throughout the nucleus, from the basal region near the tail attachment point to the apex and partially extending into the hook. Some examples of these positions in individual nuclei are shown in Figure 5.

**Figure 5.**
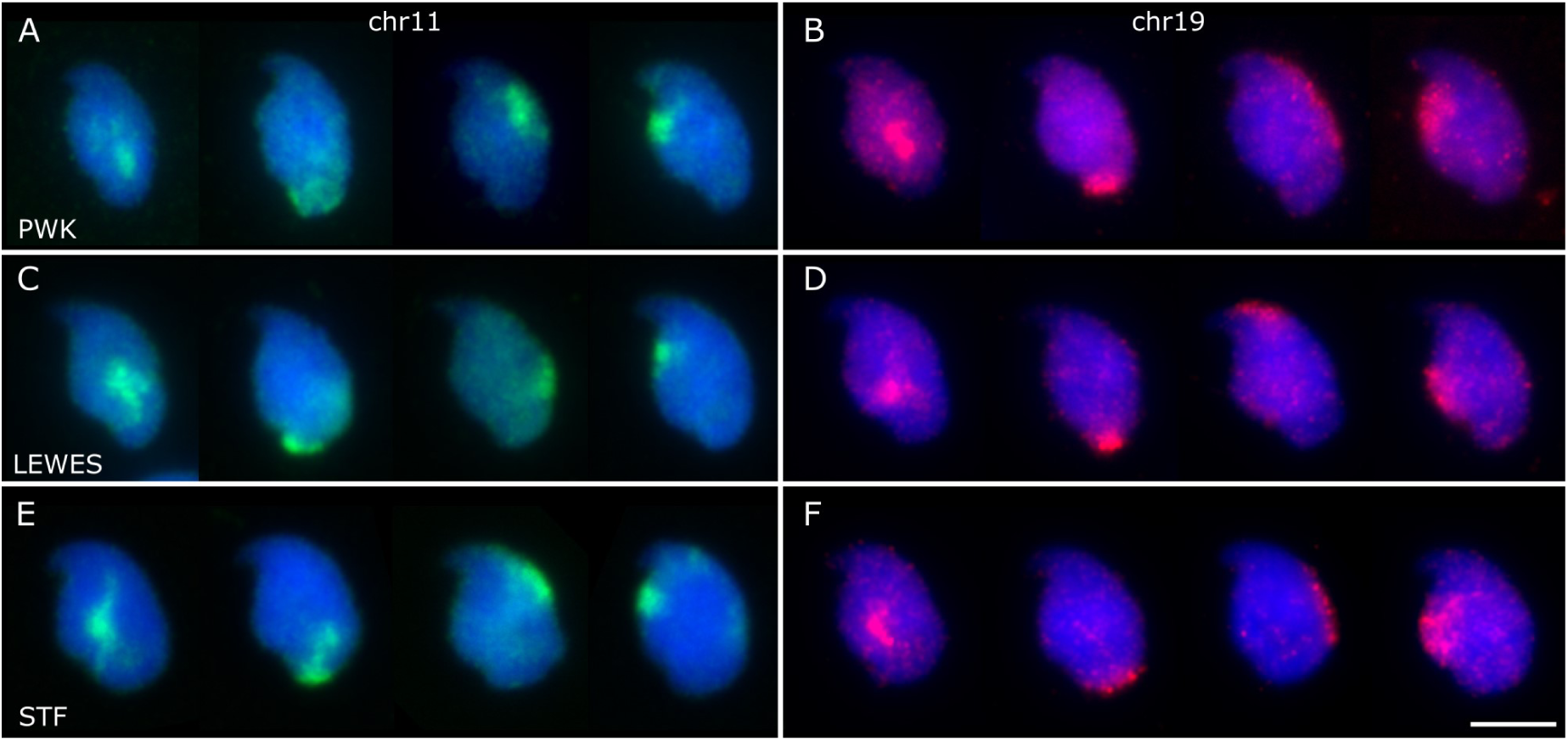
Examples of individual chromosome positions for chr11 (A, C, E) and chr19 (B, D, F) in the three strains. While the majority of the signals for each chromosome were observed ventral and basal of the nuclear centre (column 1), we found territories at the base of the nucleus (column 2), under the acrosome (column 3), and along the ventral surface below the hook (column 4). Scale bar represents 5μm.

Although hybridization efficiency was poorer in *M. spretus*, the same patterns are apparent as in the *M. musculus* strains. Interestingly, we observed instances of both chr11 and chr19 below the acrosomal curve, in which the chr19 was generally more elongated than chr11 (see Figure 5B and F). Where chromosome 19 was co-hybridised with chromosome X, we were able to see rare instances of chrX and chr19 lying adjacent, with chrX more internal (Figure S1).

Given the similarity in overall signal distributions, we looked to see if chr11 and chr19 tend to lie adjacent to each other in individual nuclei. Visually, we can see that they are occasionally adjacent, but are not always associated. Measurement of the distance between the chromosome signal centers of mass showed no difference between chr11 and 19 or between chr11 and X, nor did a comparison of warped signal images via MS-SSIM* (p>0.05, Wilcoxon rank sum tests; Figure 6). We conclude that while chromosome territories 11 and 19 may lie adjacent to each other within each individual sperm head, they are in general no closer to each other than chromosomes 11 and X. It is however important to appreciate that our data addresses chromosome territories as a whole, rather than individual loci, and further work will be needed to robustly compare our data with the Hi-C data from [17] (see also Discussion).

**Figure 6.**
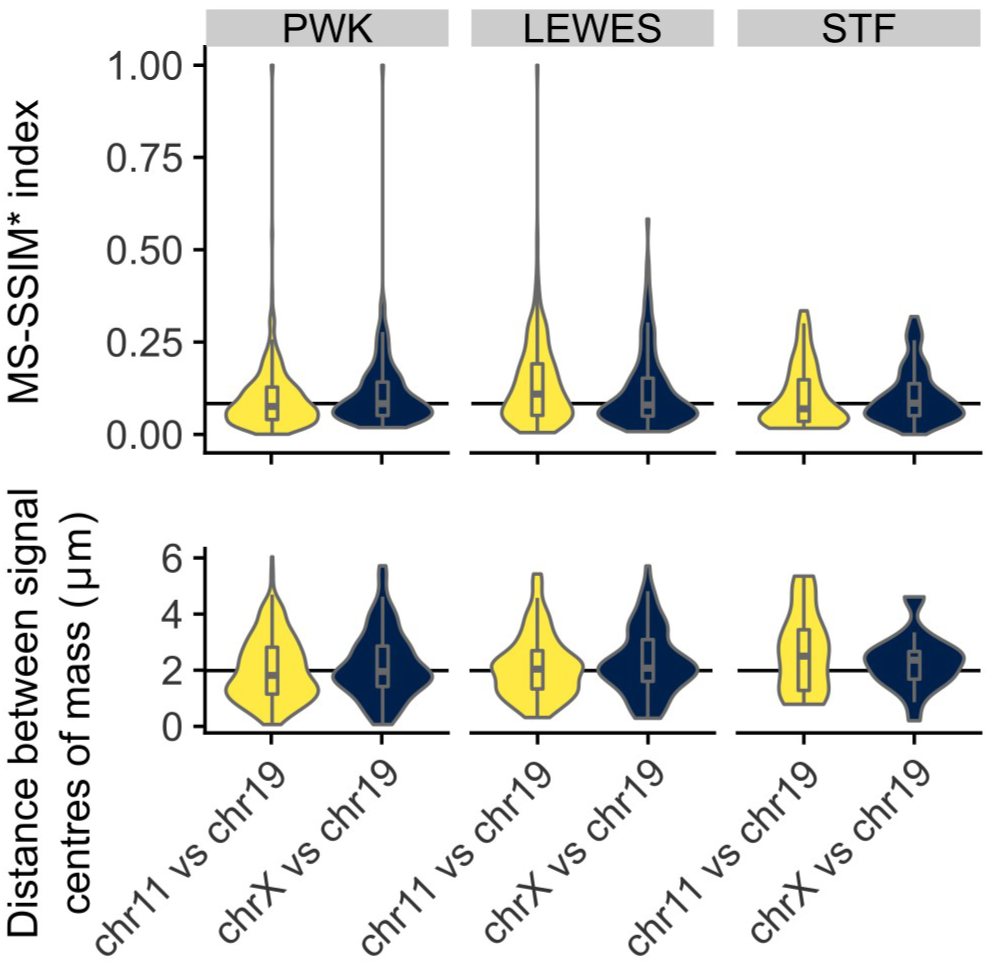
Chromosomes 11 and 19 do not colocalize within individual nuclei; colocalization of signals shows no difference comparing chr11 and chr19 as when comparing chrX and chr19 by either MS-SSIM* (upper) or the distances between the chromosome signal centers (lower).

### Quantification of signal positions reveals conserved chromosome organisation across species

In order to quantify the similarity of signal locations both within and between strains, we warped images from all three strains onto the LEWES (domesticus) consensus outline. These warped images were compared using a multi-scale structural similarity index (MS-SSIM*), a technique also used in comparisons of radiological images [24]. The X and Y territories had high structural similarity to each other in all three strains, and had high concordance between strains (Figure 7A). Similarly, we saw greater similarity between chr11 and chr19 in all three strains. The pattern was slightly less clear between *M. spretus* and the other strains, presumably due to the lower hybridisation efficiency of the probes. To confirm there was no artefactual bias introduced by the choice of LEWES as the “destination” shape, we examined the effect of warping signals onto either the PWK or STF consensus outlines, and found that this made little difference in the values obtained (see also Figure S2, Table S2). This demonstrates that our method is robust for comparing differently shaped nuclei as long as we can define structurally equivalent landmarks.

**Figure 7.**
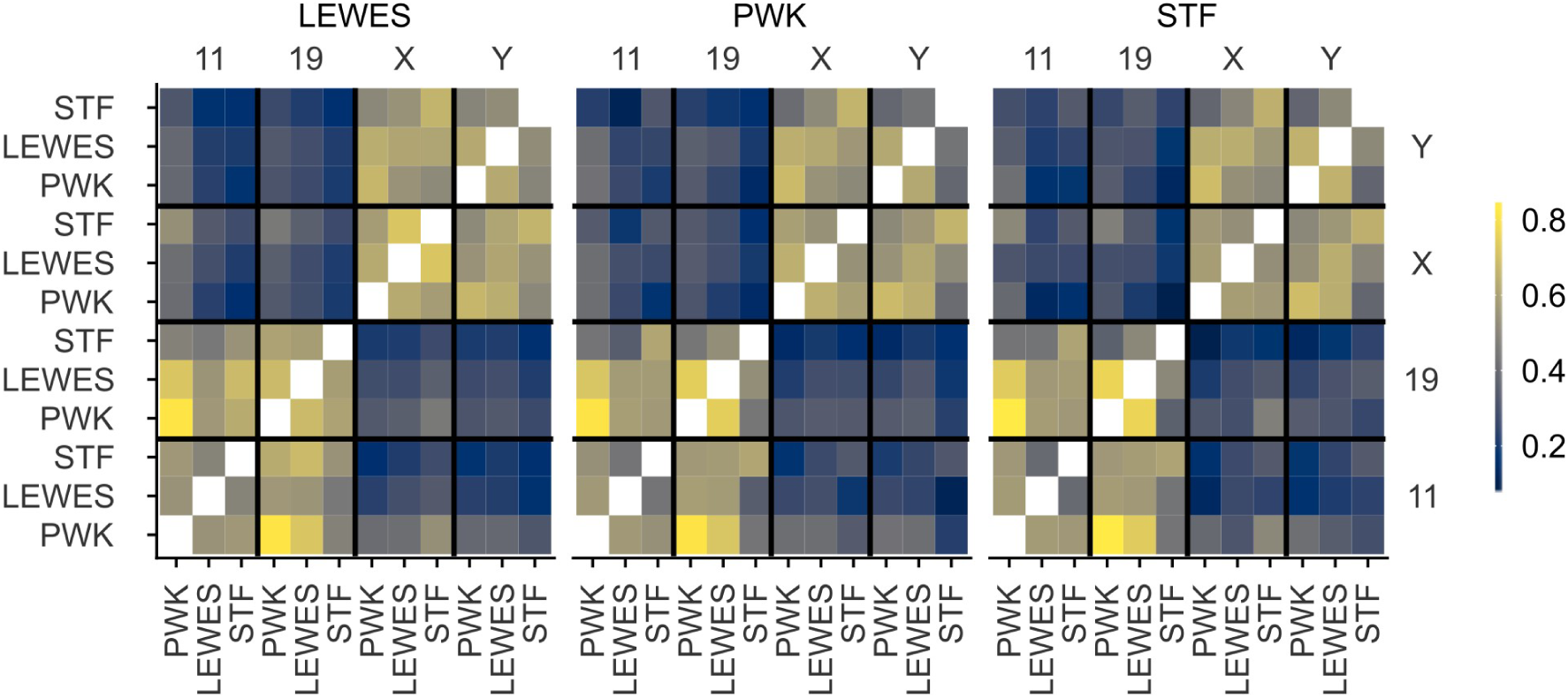
Similarity of signal distributions in composite warped images measured by MS-SSIM*, on a scale of 0-1. Images were warped in turn onto the consensus shapes of LEWES, PWK and STF. There is high correlation between the MS-SSIM* scores obtained when images are warped onto different target shapes (see Figure S2). Both within strains and between strains, there is a clear similarity between the distributions of chrX and chrY, and chr11 and chr19, but little similarity between the reciprocal combinations.

## Discussion

We have presented here a new method for quickly and efficiently mapping chromosome position in asymmetric nuclei, such as sperm, based on linking chromosome signals with morphometric information about nuclear structure. Using this analysis, we have been able to measure and quantify differences in chromosome territory position in sperm from three mouse strains. All mouse strains studied here diverged, at most, 3 million years ago [25,26], and the karyotypes of *M. musculus* and *M. spretus* both have 40 chromosomes [27]. M. *musculus* and *M. spretus* are able to produce hybrids in laboratory conditions, of which the female F1 is fertile [28]. We have demonstrated here that orthologous chromosomes adopt similar conformations in the three strains, despite differences in nuclear shape.

### Chromosomes X and Y have a conserved dorsal/sub-acrosomal position

Both the mouse X and Y chromosomes have been subject to massive amplification of euchromatic sequences. The full sequence of a M. *M. musculus* C57Bl/6 Y chromosome revealed the complex ampliconic structure [29], and demonstrated the presence of similar amplicons on the *M. spretus* Y. These amplicons are thought to arise from genomic conflict in spermatids [30], and copy number measurements of individual ampliconic genes suggests *M. spretus* has generally amplified the same gene families as *M. musculus*, with the exception of X-linked H2al1, which has amplified specifically in the *M. musculus* lineage.

Despite the close evolutionary relationship of *M. musculus* and *M. spretus*, some small rearrangements involving the sex chromosomes have been documented - for example, the Clcn4 gene, X-linked in most mammals including *M. spretus*, is autosomal in *M. musculus* [31], with clear translocation breakpoints surrounding the gene [32].

Given the overall structural similarity of the orthologous chromosomes, it is likely they occupy a similar volume within the nucleus, and are subject to similar conformational constraints. The sex chromosomes have been previously described to adopt a dorsal position in the rodent sperm nucleus [8,9], and have been seen to be sub-acrosomal in human, marsupial and monotreme sperm [14]. It has been suggested that the X chromosome - in X-bearing sperm - is the first to enter the egg during fertilisation. The position of the Y in marsupials is not reported, but as in mice, it is likely that the Y adopts the same position as X simply because the space is available. In monotremes, the platypus Y chromosomes do show a similar distribution to the X chromosomes [33]. Since the sex chromosomes are different sizes - approximately 90Mb versus 170Mb - there must be differences in the chromatin packing to allow them to occupy the same nuclear volume. In future we will be interested to study the impact of chromosome constitution on nuclear morphology.

### Chromosomes 11 and 19 have a conserved ventral/basal distribution

Chromosome 19 has been observed by others to lie in the basal region of the nucleus in approximately % of nuclei based on imaging and manually scoring at least 350 C57Bl/6 sperm nuclei [8,9]. Our results support these data, and demonstrate conservation of this position across species. The signal in *M. spretus* is less clear, likely due to the cross-species hybridisation, but the pattern is still distinguishable.

Our data from co-hybridisations suggest that although chr11 and chr19 adopt a similar overall location, they do not always lie adjacent within a single nucleus. This indicates that while they have preferred regions of the nucleus, they are mostly unconstrained with regard to each other. Aggregate data from Hi-C experiments in C57Bl/6 sperm [17] have indicated that chr19 is infrequently associated with the X chromosome (and by inference, the Y chromosome), and that chr11 is only moderately associated with both chrX and chr19. It is however currently unclear why Hi-C shows chromosome 19 to be more strongly associated with chromosome 11 than the X chromosome, given our data showing that these three chromosome territories are on average equidistant. One potential explanation is that while our measurements focus on the centre of each chromosome territory, interactions occur at the periphery of territories in cells where they abut each other. Also worthy of note is that the mouse sperm head tends to have a DAPI-dense chromocenter “core”, and that the X/Y and 11/19 regions are deduced to usually lie on opposite sides of this. Potentially this core forms a barrier to inter-chromosomal interactions. A higher resolution investigation of individual loci found to be associated in the Hi-C data will help resolve this question.

Overall, our measurements tend to support previous Hi-C and FISH findings in laboratory mouse sperm, and provide evidence that the same patterns will be found in *M. spretus*. The concept of ‘spatial synteny’ - the conserved 3D position of orthologous loci despite karyotypic rearrangements - has been proposed [34], and there is increasing evidence for conserved nuclear organization of genes following chromosomal rearrangements [35]. As we extend our studies, it will be interesting to compare the positions of the full set of chromosomes, to better understand how the shorter and fatter *M. spretus* nucleus maps on the longer, thinner *M. musculus* nucleus. Further comparisons with other mouse strains with greater shape variability will also be of value; for example BALB/c, which have frequent shape abnormalities and aneuploidies [18,36].

Studies of strains with other aneuploidies, chromosomal rearrangements or Robertsonian fusions, which will additionally constrain chromosome territories will be of interest. In humans, no gross morphological differences in sperm nuclei have been seen in men carrying Robertsonian fusions [37]. However, in boars *(Sus scrofa)*, while gross nuclear morphology was not perturbed in animals carrying a t(13;17) Robertsonian translocation, differences were apparent in the positions of the affected chromosomes [38]. Extending beyond mice, rats *(Rattus rattus)* have a much thinner hooked sperm nucleus; rat chromosomes have been mapped in developing spermatids from stages 7-13. The nucleus is compressed from a structure which at stage 10 is markedly similar to a mature mouse sperm nucleus [39]. The associated dynamics of nuclear reshaping during spermiogenesis, and chromosome repositioning are an area of active research [10].

### This method allows rapid screening of large numbers of nuclei

In this analysis, we examined more than 3000 nuclei, and the method scales to greater numbers with little additional time or user effort after images have been captured. Importantly, our analysis does not rely on extensive manual classification of chromosome position, making it less subjective than current approaches, and amenable to automation. The use of a mesh to warp signals from different nuclei onto a single template shape allows for quantitative measurements of the similarity of signal distributions between images, and in principle will allow us to study small differences in locus position that have been beyond the scope of current scoring systems. Beyond chromosome territory positioning, it is also amenable to the study of single BAC probes; together with Hi-C data this will allow us to study which intra- and inter-chromosomal folding contacts are retained in the sperm head, and address long standing questions of whether sperm chromatin organisation represents an echo of round spermatid chromatin organisation, or prefigures future chromatin folding dynamics in the fertilised zygote.

A further methodological interest would be to identify reliable internal structural features within the nucleus, using DAPI or other stains. Currently we use only peripheral features as landmarks, which puts limits on the accuracy of our mesh when deforming images. More internal structural data would permit more complex morphometric approaches such as Teichmüller mapping, which has been used successfully in analysis (for example) of wing shape in Drosophila species [40].

## Conclusions

Here we have demonstrated a new method for locating chromosome paints or other nuclear signals within mouse sperm nuclei, which is in principle also applicable to other asymmetric nuclei, including nuclei with fewer axes of asymmetry, such as spatulate sperm nuclei. We have used this technique to confirm the non-random positioning of the sex chromosomes, and of chromosomes 11 and 19, and demonstrated quantitation of signal positions allowing comparison between different strains and species. Importantly, we have integrated this method into existing open-source image analysis software designed for other biologists.

## Supplementary Materials

Figure S1: Chromosomes X and 19 co-hybridization; Figure S2: Comparison of MS-SSIM* scores using different warping templates; Table S1: Numbers of nuclei analyzed, Table S2: Complete MS-SSIM* comparisons between warped composite images.

## Author Contributions

Conceptualization, BMS and PE; Methodology, BMS and PE; Software and Validation, BMS; Investigation, JB, CCR; Data Curation and Formal Analysis, BMS; Visualization, BMS; Supervision and Project Administration, PE; Writing - Original Draft, BMS and PE; Writing - Review and Editing, BMS, CCR, JMG, ELL and PE; Resources, JMG, ELL, EEKK, NA and PE; Funding Acquisition, NA and PE.

## Acknowledgments

We thank the animal handling staff at the University of Montana.

