## Supplementary figures and images for "Automated nuclear cartography reveals conserved sperm chromosome territory localization across 2 million years of mouse evolution"

### Figure S1.tif

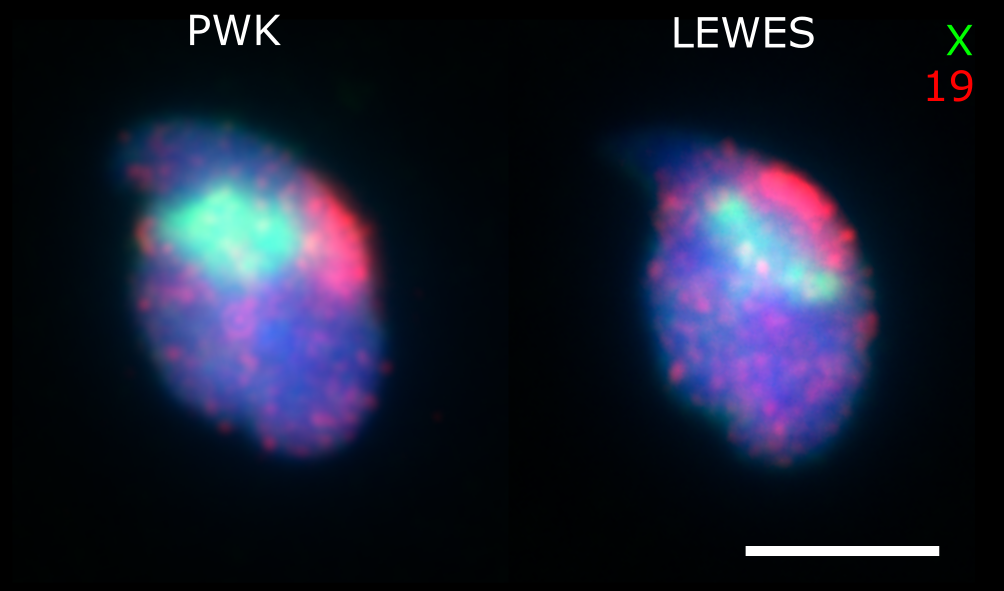

### Figure S2.tif

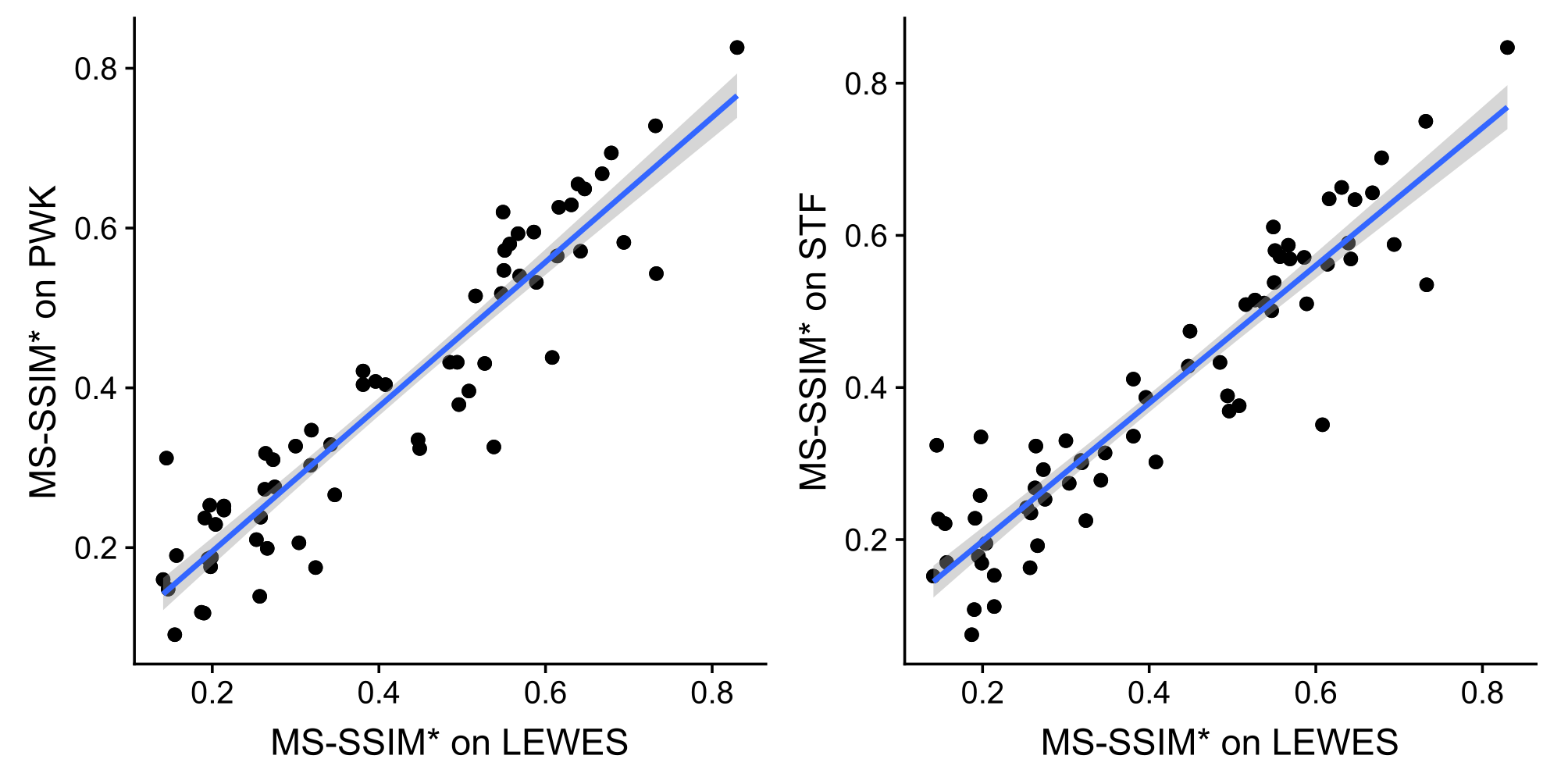
